# PTBP1 is upregulated by Zika virus infection *via* HIF-1α signal and hijacks NS1 protein to induce NS1 degradation to restrain viral replication

**DOI:** 10.1101/2024.11.19.624259

**Authors:** Menglan Rao, Zhiwei Lei, Shuang Liu, Jiuxiu Lin, Yue Kong, Yicong Liang, Zhen Luo

**Affiliations:** Department of Immunology and Microbiology, College of Life Science and Technology, Jinan University, Guangzhou 510632, China; Department of Gastroenterology, Affiliated Qingyuan Hospital, Guangzhou Medical University, Qingyuan People’s Hospital, Qingyuan 511500, China; Department of Microbiology and Immunology, Basic Medicine College, Jinan University, Guangzhou 510632, China; Institute of Medical Microbiology, Jinan University, Guangzhou 510632, China; Key Laboratory of Viral Pathogenesis & Infection Prevention and Control (Jinan University), Ministry of Education, Guangzhou 510632, China

**Keywords:** Zika virus, Polypyrimidine tract-binding protein 1, Non-structural Protein 1, Hypoxia-inducible factor-1α, lysosomal pathway

## Abstract

Zika virus (ZIKV), belonging to the *Flaviviridae* family, has been a severe threat to human health since the worldwide outbreak. ZIKV is capable of inducing fetal microcephaly, Guillain-Barré syndrome, and other serious neurological complications. Polypyrimidine tract-binding protein 1 (PTBP1) is a key member of the heterogeneous nuclear ribonucleoproteins (hnRNPs) family, functioning in selective mRNA splicing and gene expression regulation. Our previous study has indicated that the expression of PTBP1 increases in astrocytes upon ZIKV infection, yet the precise regulatory mechanisms underlying its role in viral replication remain elusive. In this study, we elucidated the specific pathway by which ZIKV upregulates PTBP1 expression through the activation of Hypoxia-inducible factor-1α (HIF-1α) expression. Further investigation revealed that overexpression of PTBP1 effectively inhibits viral replication, whereas knockdown of PTBP1 significantly enhances viral replication. Mechanistically, using co-immunoprecipitation assays for protein interaction screening, we identified an interaction between PTBP1 and ZIKV non-structural protein NS1. Detailed studies demonstrated that PTBP1 bound and colocalized with NS1 to lead to the degradation of NS1 protein via a lysosomal pathway. Collectively, our findings unveil a novel mechanism underlying that ZIKV infection induces the expression of PTBP1 via the HIF-1α pathway, subsequently the accumulated PTBP1 binds to ZIKV NS1 protein to promote NS1 degradation, thereby effectively inhibiting viral replication. The study illustrates a distinct restricted cellular factor that regulates ZIKV replication, which provides a potential target for the control of the viral replication and pathogenesis during the ZIKV epidemic.

**Importance:** Since the outbreak of ZIKV infection among human in 2014, a Zika epidemic has caused Zika fever accompanied with fetal microcephaly, Guillain-Barré syndrome, and other neurological symptoms. Emerging evidence reveals that ZIKV infects astrocytes to specially induce the expression of polypyrimidine tract-binding protein 1 (PTBP1), one of hnRNPs members. However, the interplay between PTBP1 and ZIKV replication is highly concerned. Here, we uncover a distinct manner that ZIKV infection induces PTBP1 expression through the activation of hypoxia-inducible factor-1α (HIF-1α) signal. Additionally, activation of HIF-1α signal hinders ZIKV replication relying on PTBP1 accumulation. Further investigations suggest that PTBP1 restrains ZIKV replication by interacting with ZIKV NS1 protein, thereby leading to the degradation of NS1 protein via a lysosomal pathway. Collectively, our findings illustrate a novel restricted cellular factor PTBP1 mediated by HIF-1α that regulates ZIKV replication, which provides a potential therapeutic target of the viral replication and pathogenesis against ZIKV pandemic.

## Introduction

ZIKV is an arthropod-borne single-stranded positive-sense RNA virus belonging to the Flavivirus genus within the *Flaviviridae* family (1). Although most individuals infected with ZIKV do not exhibit overt symptoms, approximately 20-25% of infected patients typically present with a series of mild clinical manifestations such as headache, fever, rash, and musculoskeletal pain (2). However, the severity of ZIKV infection lies in its potential to cause neurological complications, including fetal microcephaly and neurological defects (3), Guillain-Barre syndrome (4), meningitis, and myelitis (5). Currently, there are no specific vaccines or antiviral drugs available for the prevention or treatment of ZIKV infection, and supportive and symptomatic treatments are the mainstay for managing ZIKV-infected patients (6, 7). It is urgent to discover the exact manner that regulates ZIKV replication in the host.

The ZIKV particle is spherical with a diameter of 40-60 nm and a genome size of approximately 10.8 kb, which comprises 5’ and 3’ untranslated regions (UTRs) and a single open reading frame (ORF) (8). The ORF encodes a polyprotein that is cleaved into three structural proteins: C, prM, E, and seven non-structural proteins (NS1, NS2A, NS2B, NS3, NS4A, NS4B, and NS5) (9). Among them, the non-structural proteins play pivotal roles in virus replication, assembly, and evasion of host defenses. The NS1 protein is multifunctional and involved in virus replication and immune evasion (10). It exists in cells as monomers, dimers on the endoplasmic reticulum membrane, and hexamers outside the cell (11). The dimers participate in the formation of virus replication complexes, while the hexamers are involved in immune evasion. ZIKV NS1 and NS4B interact with TANK-binding kinase 1 (TBK1) and inhibit its phosphorylation, thereby suppressing the production of interferon-β (IFN-β) (12). ZIKV NS1 also recruits the host deubiquitinase Ubiquitin-specific protease 8 (USP8) to inhibit the degradation of caspase-1, enhancing NOD-, LRR- and pyrin domain-containing protein 3 (NLRP3) inflammasome activation and reducing type I IFN production, thus helping ZIKV evade the host antiviral response (13). Additionally, NS1 serves as an important biomarker for early diagnosis of ZIKV infection (14). Thus, the role of ZIKV NS1 in the control of viral replication is highly concerned.

Polypyrimidine Tract-Binding Protein 1 (PTBP1), also known as PTB or heterogeneous nuclear ribonucleoprotein I (hnRNP I), is a member of the hnRNP family of splicing proteins. With a relative molecular mass of 57 kDa, it is a shuttling protein primarily located in the nucleus (15). The protein structure consists of four non-canonical RNA recognition motifs, a nuclear localization domain, and a nuclear export signal at the N-terminus (16, 17). PTBP1 participates in RNA maturation, transport, localization, and translation processes, primarily serving as a splicing factor in the alternative splicing of pre-mRNA (18–20). Due to its functional diversity, it has become one of the important targets for studying gene expression regulation mechanisms and disease pathogenesis. PTBP1 is also associated with biological features under infectious conditions. Our previous study suggests the upregulation of PTBP1 companies with ZIKV infection in primary mouse astrocytes in the host transcriptome analysis (21), but the potential regulation of PTBP1 on ZIKV replication remains elusive.

In this study, we explore the precise mechanism of PTBP1 in the regulation of ZIKV replication. We found that ZIKV infection activates HIF-1α pathway to induce PTBP1 expression, in turn, the accumulation of PTBP1 binds to ZIKV NS1 protein to promote NS1 degradation, resulting in inhibiting viral replication. The study illustrates that PTBP1 regulates ZIKV replication as a distinct restricted cellular factor in the host and provides a potential target in the interventions of the ZIKV replication and pathogenesis.

## Materials and methods

### Cell lines and virus

Mosquito (C6/36) cells, African green monkey kidney (Vero) cells, human lung adenocarcinoma (A549) cells, human embryonic kidney 293T (HEK293T) cells were purchased from the American Type Culture Collection (ATCC) (Manassas, VA, USA). The cell culture was prepared with Dulbecco modified Eagle medium (DMEM) (Invitrogen; Carlsbad, CA, USA), 10% fetal bovine serum (FBS) (Gibco; Grand Island, NY, USA), 100 U/mL penicillin, and 100 mg/mL streptomycin (Gibco). All cells were cultured in a humidified incubator at 37 ℃ and 5% CO_2_. ZIKV has been previously described as an Asian strain isolate zl6006 (GenBank accession number KU955589.1) (22), obtained from the Institute of Pathogenic Microorganisms of the Guangdong Provincial Center for Disease Control and Prevention. ZIKV was propagated in C6/36 cells and the viral titer was measured in Vero cells.

### ZIKV propagation and titration

C6/36 cells were infected with ZIKV stock solution to allow the virus to incubate the cells for 2 h and then were replaced with fresh DMEM for 6-7 days until the cells caused an obvious cytopathic effect. The cell supernatants were collected and centrifuged at 1000 × *g* for 3 min. The supernatant was filtered using a 0.22 μm filter to obtain the virus stock solution. The TCID50 assay was used to calculate the ZIKV titer. Vero cells were incubated with ZIKV at an appropriate multiplicity of infection (MOI) for 2 h. The culture medium was replaced with the fresh medium to continue culture for 3-5 days. After reaching the predetermined time point, the cells were observed for subsequent analysis. The effect of viral infection on the control group was compared and evaluated. Reed Muench method was used to calculate the plaque forming unit (PFU) of the virus.

### Antibodies, Reagents, and Plasmids

Mouse monoclonal antibody against GAPDH (Cat: G9295) was purchased from Gibco. Rabbit polyclonal antibodies against PTBP1 (Cat:12582-1-AP) and HIF-1α (Cat: 20960-1-AP), rabbit IgG control polyclonal antibody (Cat: 30000-0-AP), rabbit monoclonal antibody against HA (Cat: 51064-2-AP), HRP-conjugated Affinipure goat anti-mouse IgG (Cat: SA00001-1) and goat anti-mouse IgG (Cat: SA00001-2) were obtained from Proteintech Group (Chicago, IL, USA). Rabbit polyclonal antibodies against ZIKV NS5 (Cat: GTX133312) and NS1 (Cat: GTX634159) were obtained from Genetex Inc. (Irvine, CA, USA). Mouse monoclonal antibody against FLAG (Cat: F3165) was purchased from Sigma (St Louis, MO, USA). FITC-conjugated Affinipure donkey anti-rabbit IgG (Cat: SA00003-8) and CoraLite594-conjugated donkey anti-mouse IgG (Cat: SA00013-7) were purchased from Proteintech. IOX2 (Cat: HY-15468), YC-1 (Cat: HY-14927), 3-MA (Cat: HY-19312), MG132 (Cat: HY-13259) and NH_4_Cl (Cat: HY-Y1269C) were purchased from MedChemExpress (Princeton, NJ, USA). LysoTracker Deep Red (Cat: L12492) was purchase from Invitrogen (Carlsbad, CA, USA). DAPI (Cat: 10236276001) was purchased from Roche (Basel, Switzerland).

The cDNA encoding human PTBP1 cloned into pCAGGS-HA vector was constructed by a standard molecular cloning method. The plasmids encoding ZIKV non-structural proteins, FLAG-NS1, NS3, NS4A, NS4B, NS5, or pcDNA3.1 (+)-3 × FLAG vector were previously described (23).

### RNA extraction and qRT-PCR

Cells were lysed using Trizol reagent (Invitrogen) for 10 min at room temperature and total RNA was extracted. Then, a certain amount of RNA sample was generated to cDNA using the reverse transcription kit (Cat: RRO36A) Takara Bio Inc). The cDNA samples in triplex were subjected to qPCR detection by Light Cycler 480 (Roche Diagnostics) and SYBR Green Real-Time PCR Master Mix (Roche Di Biologists). The value was calculated by using 2^−△△CT^, and all data were normalized to GAPDH. The primers used in this study as follows:

GAPDH F: 5’-GTCTCCTCTGACTTCAACAGCG;

GAPDH R: 5’-ACCACCCTGTTGCTGTAGCCAA;

PTBP1 F: 5’-CTCCAAGTTCGGCACAGTGTTG;

PTBP1 R: 5’-CAGGCGTTGTAGATGTTCTGCC;

shPTBP1 F: 5’-CCGGGCCAACACCATGGTGAACTACCTCGAGGTAGTTCACCATGGTGTTGGCT TTTTG;

shPTBP1 R: 5’- AATTCAAAAAGCCAACACCATGGTGAACTACCTCGAGGTAGTTCACCATGGTGTT GGC;

ZIKV F: 5’-GGTCAGCGTCCTCTCTAATAAACG;

ZIKV R: 5’- GCACCCTAGTGTCCACTTTTTCC.

### Lentiviral packaging

HEK293T cells were transfected with psPAX2, pMD2.G, and lentiviral vector plasmid (pLenti or pLKO.1) containing the target gene or empty plasmid. After 48 h, the lentiviral supernatant was collected and centrifuged at 1500 × *g* for 5 min. The supernatant was filtered through a 0.45 μm filter. Lentiviral infection solution containing an appropriate amount of polyglutamine was prepared and added to the cells for 48 h of continuous infection. The cells were incubated with 0.5 μg/mL puromycin for one week until the cells were no longer dead, and then the expression efficiency was measured by Western blot.

### Western blot

Cells were lysed in RIPA buffer for 1 h at 4℃ and then centrifuged at 4 ℃, 13,000 × *g* for 10 min to collect the cell lysate. The protein concentration was measured using the BCA assay kit (Beyotime Biotechnology; Haimen, Jiangsu, China). Proteins were separated by SDS-PAGE and transferred to the PVDF membrane. The membrane was blocked with TBST containing 5% skimmed milk powder for 1 h at room temperature. After 3 times washing with TBST, the membranes were incubated with primary antibody overnight at 4 ℃. After the last wash with TBST, the membranes were incubated with secondary antibodies for 1 h. The protein bands were captured using a chemiluminescence imaging system (Bio-Rad; Hercules, CA, USA). The protein expression relative to internal control is quantified using Image J software.

### Co-immunoprecipitation assay

HEK293T cells were transfected with indicated plasmids and harvested 36 h later. The cells were lysed with RIPA lysis buffer containing protease inhibitors and then centrifuged at 12,000 × *g*, 4℃ for 10 min. The cell lysates were incubated with FLAG-Trap Protein-G Sepharose (GE Healthcare, Milwaukee, WI, USA) solution for 4 h. The pellets were washed with lysis buffer. The samples were suspended by adding 2 × loading buffer and subjected to Western blot analysis.

### Immunofluorescence assay

Cells were fixed with 4% paraformaldehyde for 30 min and washed three times with PBS for 5 min each time. Cells were permeabilized with 0.4% Triton X-100 for 20 min and washed three times with PBS for 5 min each time. Then, a 5% BSA blocking solution was added and blocked at room temperature for 30 min. After blocking, cells were incubated with primary antibodies, then fluorescent secondary antibodies and DAPI. For the staining of endogenous PTBP1 and ZIKV NS1, two rabbit antibodies were labeled green and red dye, respectively, with a multiplex immunohistochemistry (IHC)/immunofluorescence (IF) staining kit (Cat: RK05903; ABclonal Biotech, Wuhan, China) according to manufacturer’s instructions. After cells were washed and observed under confocal laser scanning microscope (Leica TCS SP8; Heidelberg, Germany).

### Statistical analysis

All data were reproducible to obtain similar results. The statistical significance of the comparison of the two means was evaluated by an unpaired Student’s *t*-test. GraphPad Prism 8 software (San Diego, CA, USA) was used for analysis. The statistical significance is expressed as follows: *, *P* < 0.05, **, *P* < 0.01, and ***, *P* < 0.001.

## Results

### ZIKV infection specifically induces PTBP1 expression in A549 cells

To investigate the effect of ZIKV infection on the expression of PTBP1 in A549 cells, we first infected the cells with ZIKV for varying durations. The results displayed that ZIKV RNA level increased from 12 to 48 h p.i. (Fig 1A), suggesting a robust viral replication in A549 cells. Meanwhile, the mRNA (Fig 1B) and protein (Fig 1C) levels of PTBP1 were unregulated from 12 to 48 h p.i. upon ZIKV infection, indicating that ZIKV induces PTBP1 expression in a time-dependent manner. Consistently, we then infected the cells with ZIKV for varying multiplicities of infection (MOIs). ZIKV RNA level increased at an MOI from 0.25 to 1.0 (Fig 1D). The mRNA (Fig 1E) and protein (Fig 1F) levels of PTBP1 were induced at an MOI from 0.25 to 1.0 upon ZIKV infection, indicating that ZIKV induces PTBP1 expression in a dose-dependent manner. These findings collectively confirm that ZIKV infection typically induces PTBP1 expression in A549 cells.

**Fig. 1.**
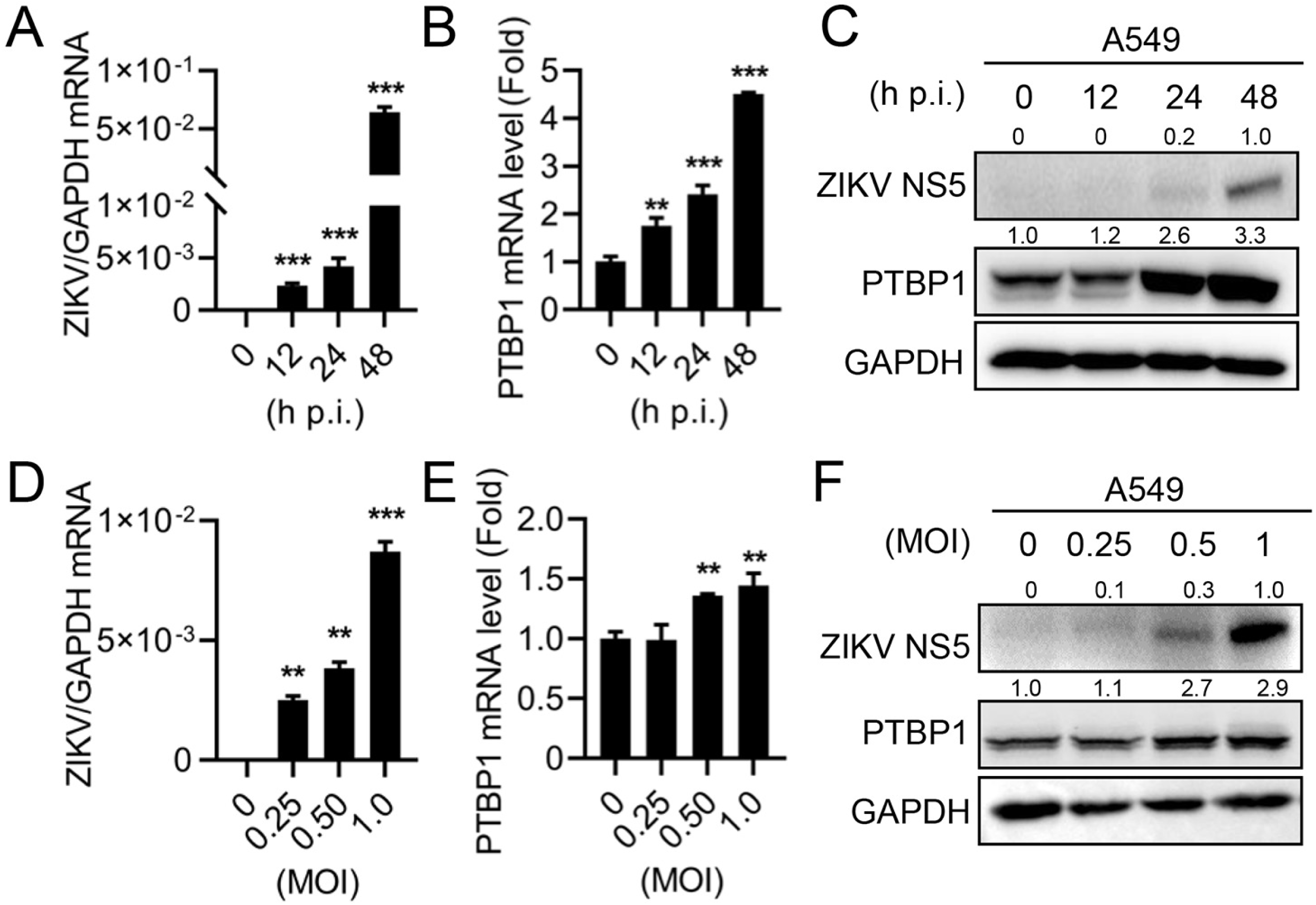
ZIKV infection upregulates PTBP1 expression in A549 cells. (A-C) A549 cells were infected with ZIKV at an MOI of 1 for different durations (0, 12, 24, and 48 h). The cells were collected for ZIKV RNA level detected by qRT-PCR (A), PTBP1 mRNA (B), and protein (C) levels detected by qRT-PCR and Western blot, respectively. (D-F) A549 cells were infected with ZIKV at MOIs of 0, 0.25, 0.5, and 1.0 for 24 h. ZIKV RNA level was detected by qRT-PCR (A), while PTBP1 mRNA (B) and protein (C) levels were detected by qRT-PCR and Western blot, respectively. Graphs were expressed as Mean±SD, n = 3. ns, not-significant; **, *P* < 0.01; ***, *P* < 0.001.

### ZIKV infection upregulates PTBP1 expression by activating the HIF-1α signal

The specific mechanism by which ZIKV upregulates the expression of PTBP1 remains unclear. Hypoxic and cellular stress responses are triggered by a process that depends on the mediation of HIF-1α (24). Under hypoxic conditions, HIF-1α can bind to the promoter of mouse PTBP1 to promote its expression (25). Based on this, we subspecialized that ZIKV upregulates PTBP1 by inducing oxidative stress to activate HIF-1α. Upon ZIKV infection in A549 cells, the expressions of both HIF-1α and PTBP1 proteins were upregulated as ZIKV replicated in a time-dependent fashion (Fig 2A), implying ZIKV infection activates the HIF-1α signal. To confirm the effect of HIF-1α activation on the PTBP1 expression, A549 cells were stimulated with an HIF-1α modulator IOX2. The expression of both PTBP1 RNA (Fig 2B) and protein (Fig 2C) increased with the growing IOX2 concentrations.

**Fig. 2.**
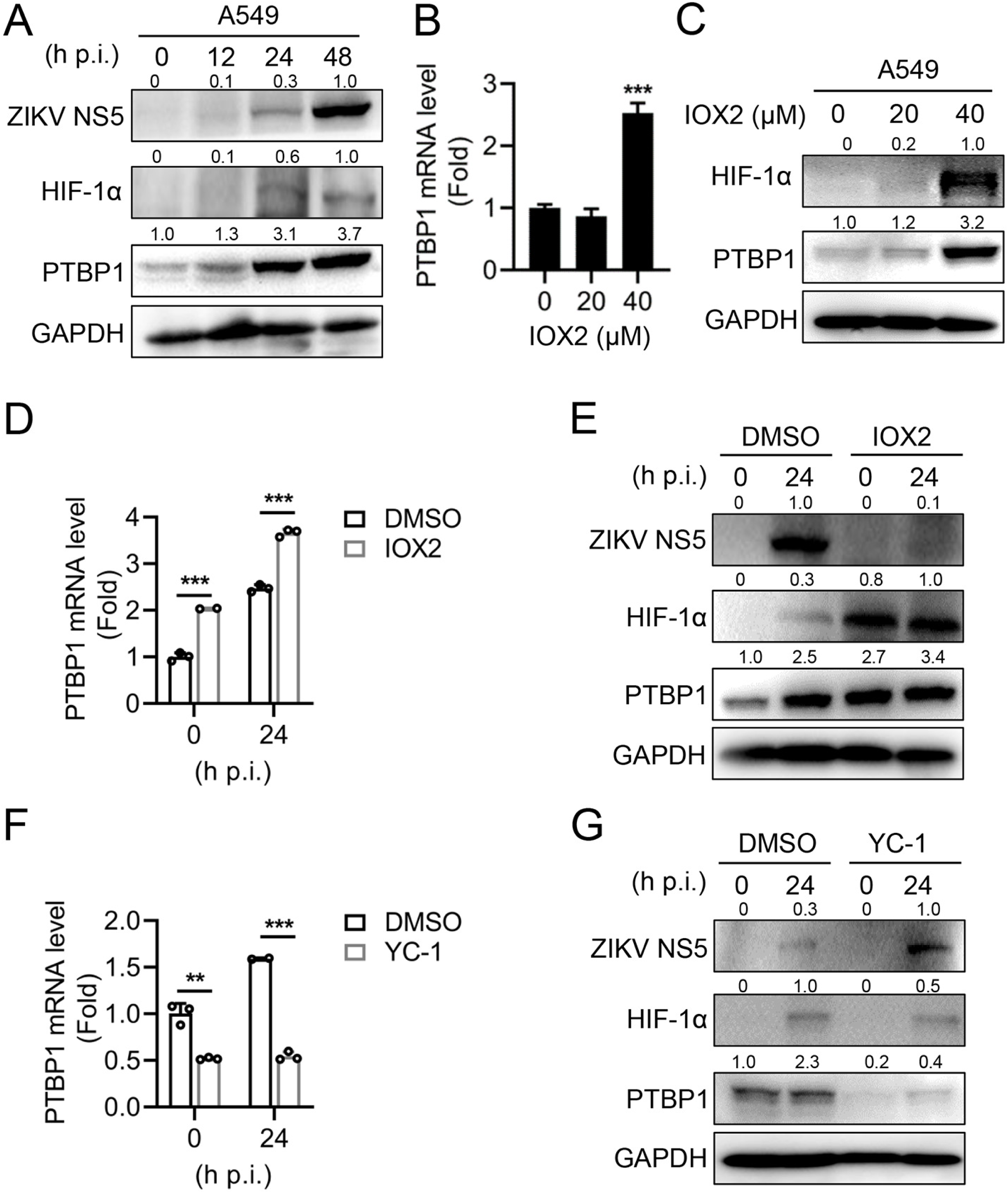
ZIKV infection promotes PTBP1 expression via HIF-1α signal. (A) A549 cells were infected with ZIKV at an MOI of 1 at different time points (0, 12, 24, and 48 h). Cells were collected at the corresponding time points for Western blot analysis. (B and C) A549 cells were stimulated with different doses of IOX2 (0, 20, and 40 μM) for 12 h. Cells were collected for qRT-PCR (B) and Western blot (C) analysis, respectively. (D and E) A549 cells were stimulated with IOX2 (40 μM) for 12 h, followed by infection with ZIKV (MOI = 1) for an additional 24 h. Cells were collected for qRT-PCR (D) and Western blot (E) analysis, respectively. (F and G) A549 cells were stimulated with a HIF-1α inhibitor YC-1 (20 μM) for 16 hours, followed by infection with ZIKV (MOI = 1) for another 24 h. Cells were collected for qRT-PCR (F) and Western blot (G) analysis, respectively. Graphs were expressed as Mean±SD, n = 3. ns, not-significant; **, *P* < 0.01; ***, *P* < 0.001.

To further investigate the role of the HIF-1α signal on the induction of PTBP1 during ZIKV infection, A549 cells were stimulated with IOX2, followed by infection with ZIKV. When HIF-1α is activated by IOX2 before ZIKV infection, HIF-1α signal enhanced ZIKV-induced PTBP1 expression in both RNA (Fig 2D) and protein (Fig 2E) levels. In contrast, when the HIF-1α signal was inhibited by an inhibitor YC-1, the upregulation of PTBP1 RNA (Fig 2F) and protein (Fig 2G) induced by ZIKV was significantly suppressed. Thus, these results support the evidence that ZIKV upregulates PTBP1 expression by activating the HIF-1α signal.

### PTBP1 inhibits ZIKV replication

Since the upregulation of PTBP1 is affected by ZIKV infection, suggesting a pivotal role of PTBP1 during ZIKV replication. To this end, we constructed a FLAG-tagged PTBP1 expressing lentivirus and successfully established an A549 cell line, LV-PTBP1, that stably expresses PTBP1, along with a control cell line, LV-Vector. The overexpression of PTBP1 was verified by Western blot analysis (Fig 3A). We observed that overexpression of PTBP1 significantly reduced the levels of ZIKV RNA (Fig 3B) and viral NS5 protein (Fig 3C). Furthermore, TCID_50_ assays confirmed that overexpression of PTBP1 inhibited the production of ZIKV progeny (Fig 3D). Conversely, we also constructed shPTBP1 RNA expressing lentivirus and established shPTBP1 cells and control cells (shNC). The knockdown efficiency was confirmed by Western blot analysis (Fig 3E). In the cells that PTBP1 was knocked-down, the levels of ZIKV RNA (Fig 3F) and viral NS5 protein (Fig 3G) increased significantly. In parallel, the production of ZIKV progeny was inhibited by shPTBP1 at 24 and 48 p.i. (Fig 3H). Therefore, the data present that PTBP1 plays an inhibitory role in ZIKV replication.

**Fig. 3.**
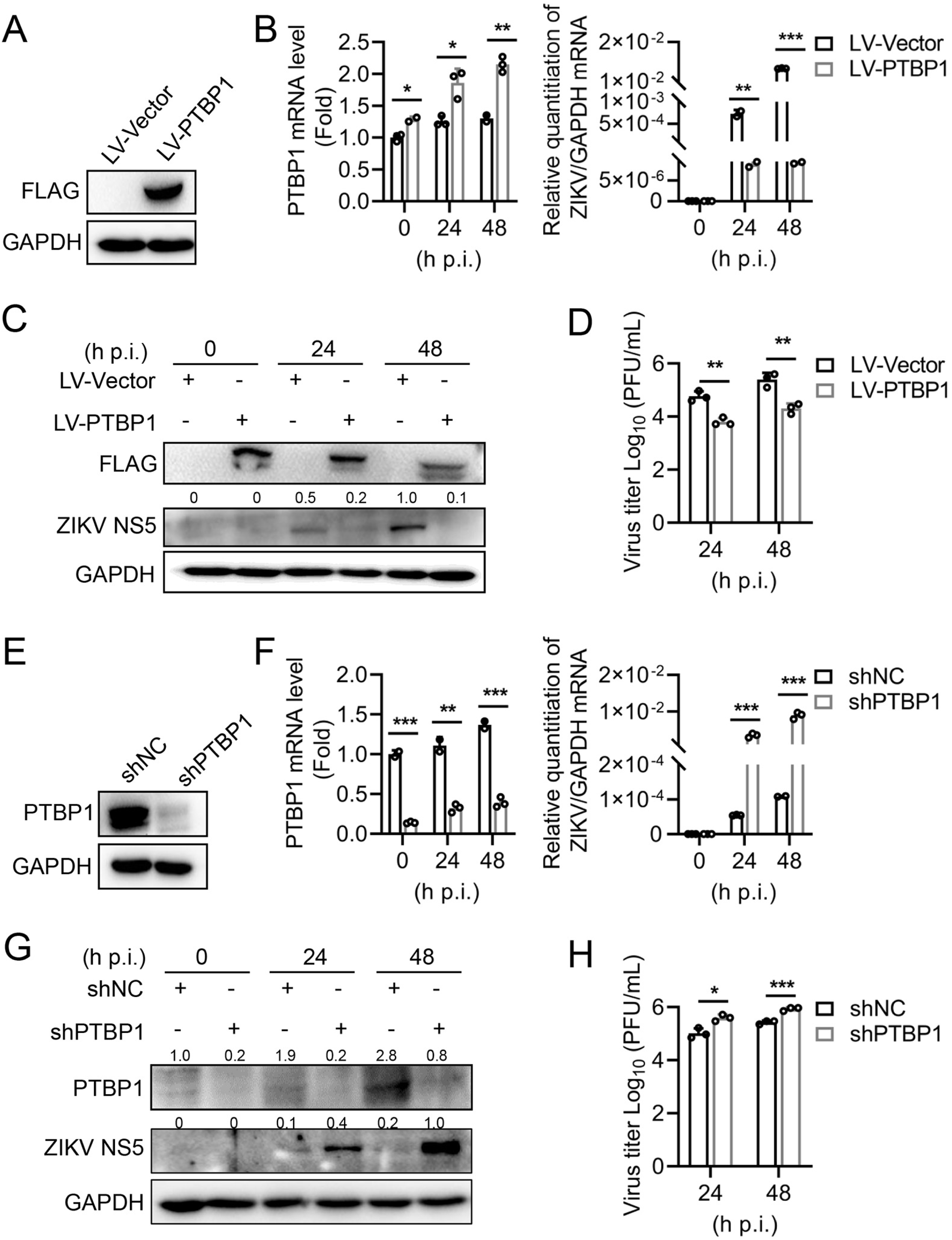
PTBP1 suppressed ZIKV replication. (A) LV-PTBP1 and LV-Vector control cells were subjected to Western blot analysis to detect the expression of FLAG-tagged protein. (B-D) LV-PTBP1 and LV-Vector cells were infected with ZIKV (MOI = 1) at different time points (0, 24, and 48 h). Cells were collected at the corresponding time points for qRT-PCR (B) and Western blot (C) analysis, respectively. Supernatants was subjected to TCID50, and the virus titer was calculated (D). (E) The stable expressing shPTBP1 and shNC control cells were subjected to Western blot analysis. (F-H) shPTBP1 and shNC cells were infected with ZIKV (MOI = 1) at different time points (0, 24, and 48 h). Cells were collected at the corresponding time points for qRT-PCR (F) and Western blot (G) analysis, respectively. Supernatants were subjected to TCID50, and the virus titer was calculated (H). Graphs were expressed as Mean±SD, n = 3. *, *P* < 0.05; **, *P* < 0.01; ***, *P* < 0.001.

### PTBP1 mediates an inhibitory role on ZIKV replication driven by HIF-1α

Considering HIF-1α activation promotes PTBP1 expression and PTBP1 inhibits ZIKV replication, we assessed the HIF-1α-driven PTBP1 upregulation whether possesses an antiviral effect on ZIKV. We pretreated A549 cells with IOX2 at varying concentrations and then infected them with ZIKV. We observed that ZIKV RNA level was decreased as the concentrations of IOX2 increased in shNC cells (Fig 4A), suggesting HIF-1α activation inhibited ZIKV replication. However, IOX2 inhibited-ZIKV replication was restored in shPTBP1 cells (Fig 4A). Parallelly, IOX2 induced HIF-1α and PTBP1 expression, resulting in the inhibition of ZIKV replication (Fig 4B), whereas such inhibitory role on ZIKV viral protein (Fig 4B) and progeny production (Fig 4C) was attenuated in the presence of shPTBP1. Therefore, PTBP1 plays an inhibitory role in ZIKV replication driven by HIF-1α.

**Fig. 4.**
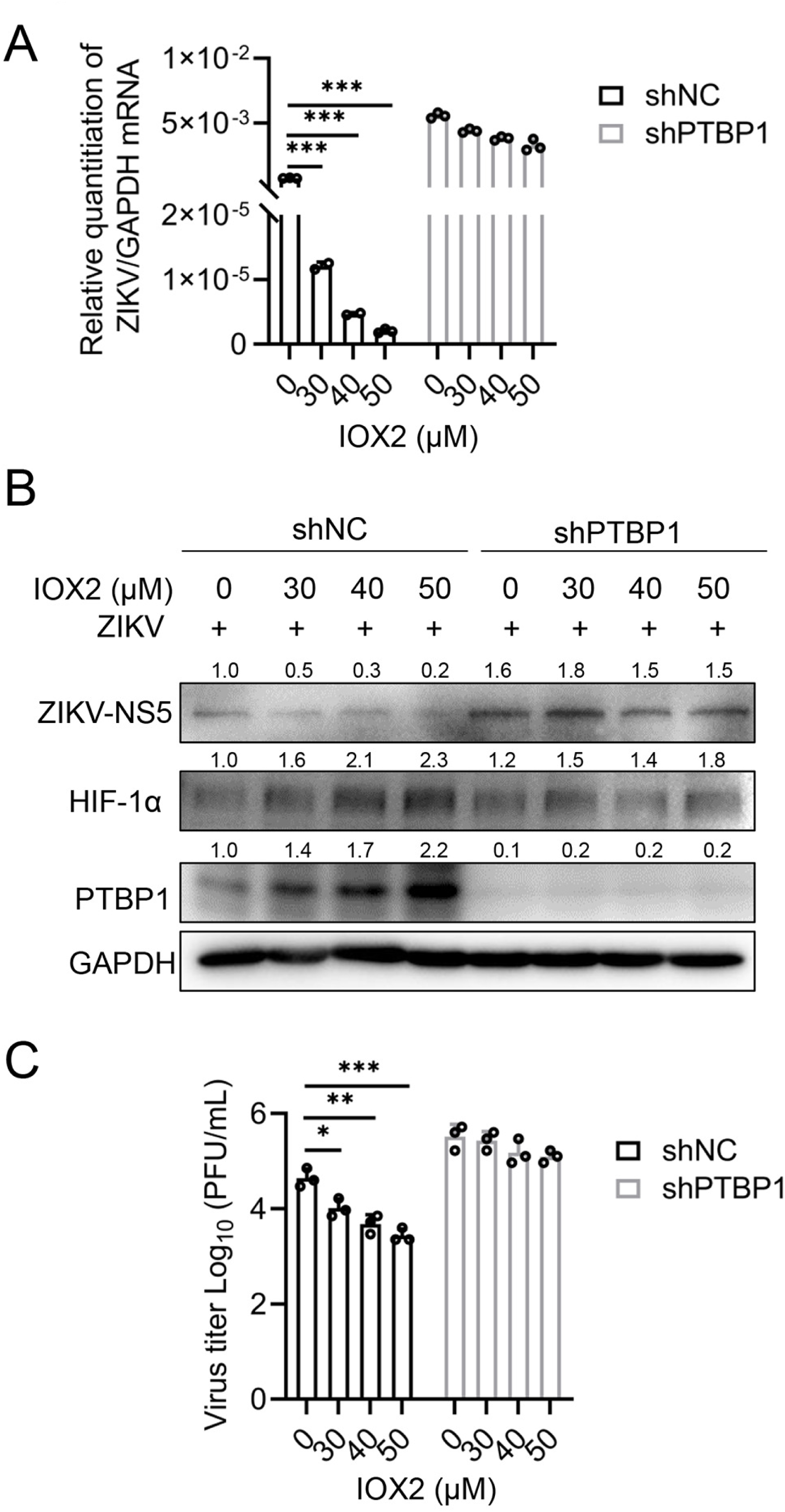
PTBP1 mediates HIF-1α-driven antiviral activity against ZIKV. (A-C) shPTBP1 cells and shNC control cells were treated with different concentrations of IOX2 (0, 30, 40, 50 μM) for 12 hours and then infected with ZIKV (MOI = 1). Cells were collected 24 hours post-infection for qRT-PCR (A) and Western blot (B) analysis, respectively. Supernatants were subjected to TCID50, and the virus titer was calculated (C). Graphs were expressed as Mean±SD, n = 3. *, *P* < 0.05; **, *P* < 0.01; ***, *P* < 0.001.

### PTBP1 interacts with ZIKV NS1

To delve into the exact mechanisms by which PTBP1 inhibits ZIKV infection, we focused on the relationship between PTBP1 and the non-structural proteins of ZIKV. We co-transfected the HA-PTBP1 plasmid with an individual plasmid encoding FLAG-tagged ZIKV non-structural proteins, including FLAG-NS1, FLAG-NS3, FLAG-NS4A, FLAG-NS4B, and FLAG-NS5, followed by conducting Co-IP experiments in HEK293T cells. The results demonstrated that PTBP1 was specifically immunoprecipitated with the ZIKV NS1 protein (Fig 4A), suggesting PTBP1 binds to viral NS1 protein. To further validate this interaction, we performed exogenous proteins by Co-IP assays in HEK293T cells after co-transfecting HA-PTBP1 and FLAG-NS1 plasmids. In pull-down assays using FLAG (Fig 4B) or HA (Fig 4C) antibodies, respectively, we affirmed the interaction between PTBP1 and ZIKV NS1.

Next, we further verified the interaction between two endogenous proteins in ZIKV-infected A549 cells. The PTBP1 antibody immunoprecipitated endogenous PTBP1 and NS1 in ZIKV-infected cells (Fig 4D). In the immunofluorescence assay, we observed the intracellular localization of PTBP1 in the nucleus in the mock-infected cells, whereas PTBP1 was mainly distributed cytoplasm in cells where the ZIKV replication occurred (Fig 4E). Importantly, the PTBP1 in cytoplasm strongly colocalized with ZIKV NS1 (Fig 4E), indicating the interaction between endogenous PTBP1 and NS1. Thus, these data reveal that PTBP1 interacts with ZIKV NS1 to inhibit viral replication.

### PTBP1 induces the degradation of ZIKV NS1 protein through the lysosomal pathway

To get a full understanding of how PTBP1 interacts with the ZIKV NS1 protein to inhibit viral replication, we investigated the impact of PTBP1 on the stability of the ZIKV NS1 protein. In the presence of the increasing amount of the HA-PTBP1, the expression level of ZIKV NS1 protein was significantly reduced, while the expression level of either ZIKV NS5 or GFP protein remained unchanged (Fig 6A), indicating that PTBP1 specifically degrades the ZIKV NS1 protein.

**Fig. 5.**
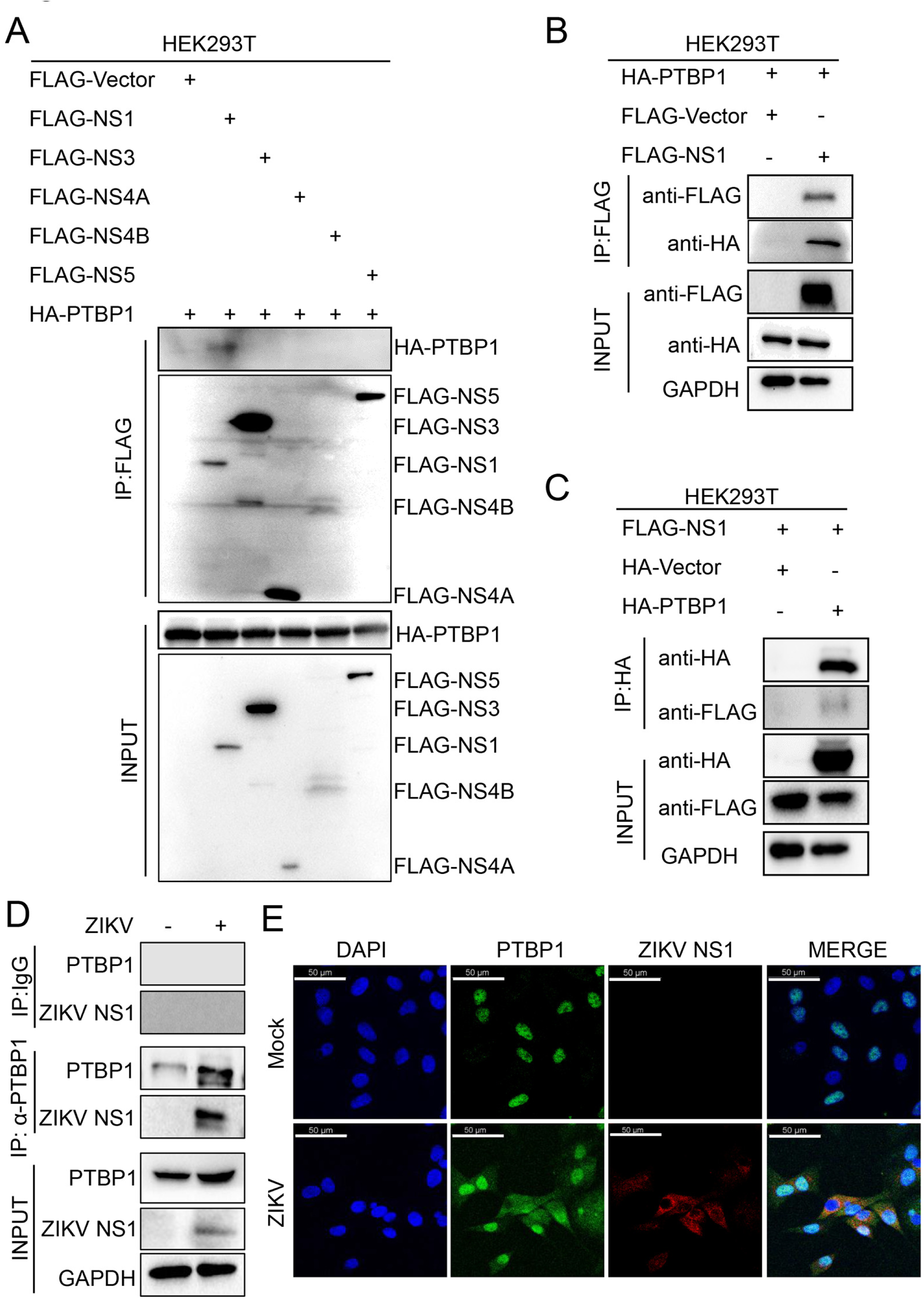
PTBP1 bind to ZIKV NS1. (A-C) HEK293T cells were transfected with the corresponding plasmids and harvested after 30 h. The cell lysates were incubated with anti-FLAG (A and B) or anti-HA (C) primary antibody overnight to immunoprecipitate target proteins, followed by Western blot analysis. (D) A549 cells were infected with ZIKV (MOI = 2) for 24 h and then harvested. The cell lysates were immunoprecipitated with anti-PTBP1 antibody, followed by Western blot analysis with indicated antibodies. (E) A549 cells were infected with ZIKV (MOI = 2) for 24 h and then incubated with primary antibodies against PTBP1 and ZIKV NS1. The treated cells were labeled with corresponding fluorescent secondary antibodies and examined under a confocal fluorescence microscope. The subcellular distribution of PTBP1 (green) and ZIKV NS1 (red) along with the nucleus stained with DAPI (blue) were captured. Scale bar = 50 μm.

**Fig. 6.**
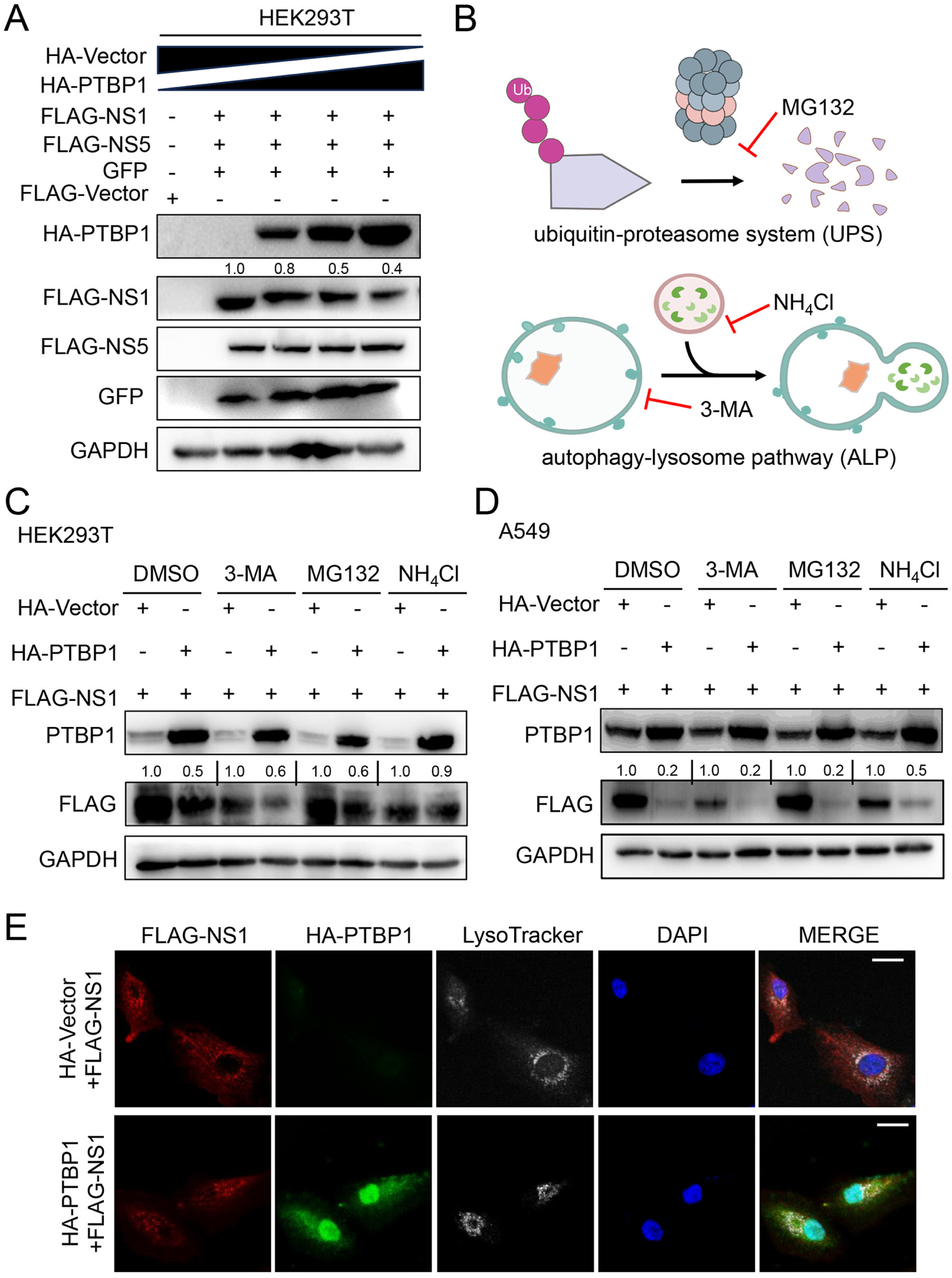
PTBP1 induces the degradation of ZIKV NS1 protein through the lysosomal pathway. (A) HEK293T cells were transfected with the HA-Vector (2.0, 1.75, 1.5, 1.25, and 0.75 μg) and FLAG-Vector (1.0, 0, 0, 0, 0 μg) or FLAG-NS1 (0, 0, 0, 0, 0.5 μg), or FLAG-NS5 (0, 0, 0, 0, 0.5 μg) or HA-PTBP1 (0, 0, 0.25, 0.5, 1.0) along with eGFP-C1 (0, 0.25, 0.25, 0.25, 0.25 μg) plasmids. After 30 h, the cells were collected and subjected to Western blot analysis. (B) The scheme of two main models of protein degradation: UPS (ubiquitin-proteasome system) and ALP (autophagy-lysosome pathway). (C and D) HEK293T cells (C or A549 cells (D) were transfected with 1.5 μg of each plasmid: HA-Vector, FLAG-NS1, and HA-PTBP1, along with FLAG-NS1. At 18 h post-transfection, the cells were treated with DMSO, 3-MA (1.0 mM), MG132 (2.5 μM), or NH_4_Cl (5 mM) for another 12 h. The protein levels were analyzed by Western blot. (E) A549 cells were transfected with 1.0 μg of each plasmid: HA-Vector or HA-PTBP1 along with FLAG-NS1. At 24 h post-transfection, the cells were labeled with anti-FLAG and anti-HA primary antibodies followed by corresponding fluorescent secondary antibodies. The lysosomes were stained with Lyso-Tracker, while nucleus was stained with DAPI. The images displaying NS1 (red), PTBP1 (green), Lysosome (white), and DAPI (blue) were captured under a confocal fluorescence microscope. Scale bar = 20 μm.

Given the protein degradation belongs to two main models: UPS (ubiquitin-proteasome system) and ALP (autophagy-lysosome pathway) (Fig 6B), we applied autophagy inhibitor 3-MA, proteasome inhibitor MG132, and lysosomal inhibitor NH_4_Cl to treat cells co-expressing PTBP1 and NS1. In HEK293T cells, PTBP1 reduced NS1 protein level, however, the NH_4_Cl restored the degradation of ZIKV NS1 protein induced by PTBP1, while 3-MA and MG132 had no such effects (Fig 6C). In A549 cells, we also observed similar results (Fig 6D), suggesting PTBP1 induced the degradation of the ZIKV NS1 protein through the lysosomal pathway. Furthermore, we observed the subcellular distribution of NS1 protein and found a certain portion of NS1 protein located in the lysosome in the absence of PTBP1 (Fig 6E). Nevertheless, the portion of NS1 protein located in the lysosome was eliminated in the presence of PTBP1 (Fig 6E). Together, these results illustrated that PTBP1 induces the degradation of ZIKV NS1 protein via the lysosomal pathway. In sum, our findings exhibited that ZIKV infection induces the expression of PTBP1 via the HIF-1α pathway, in turn, the PTBP1 binds ZIKV NS1 protein to promote NS1 degradation, thereby effectively inhibiting viral replication (Fig 7).

**Fig. 7.**
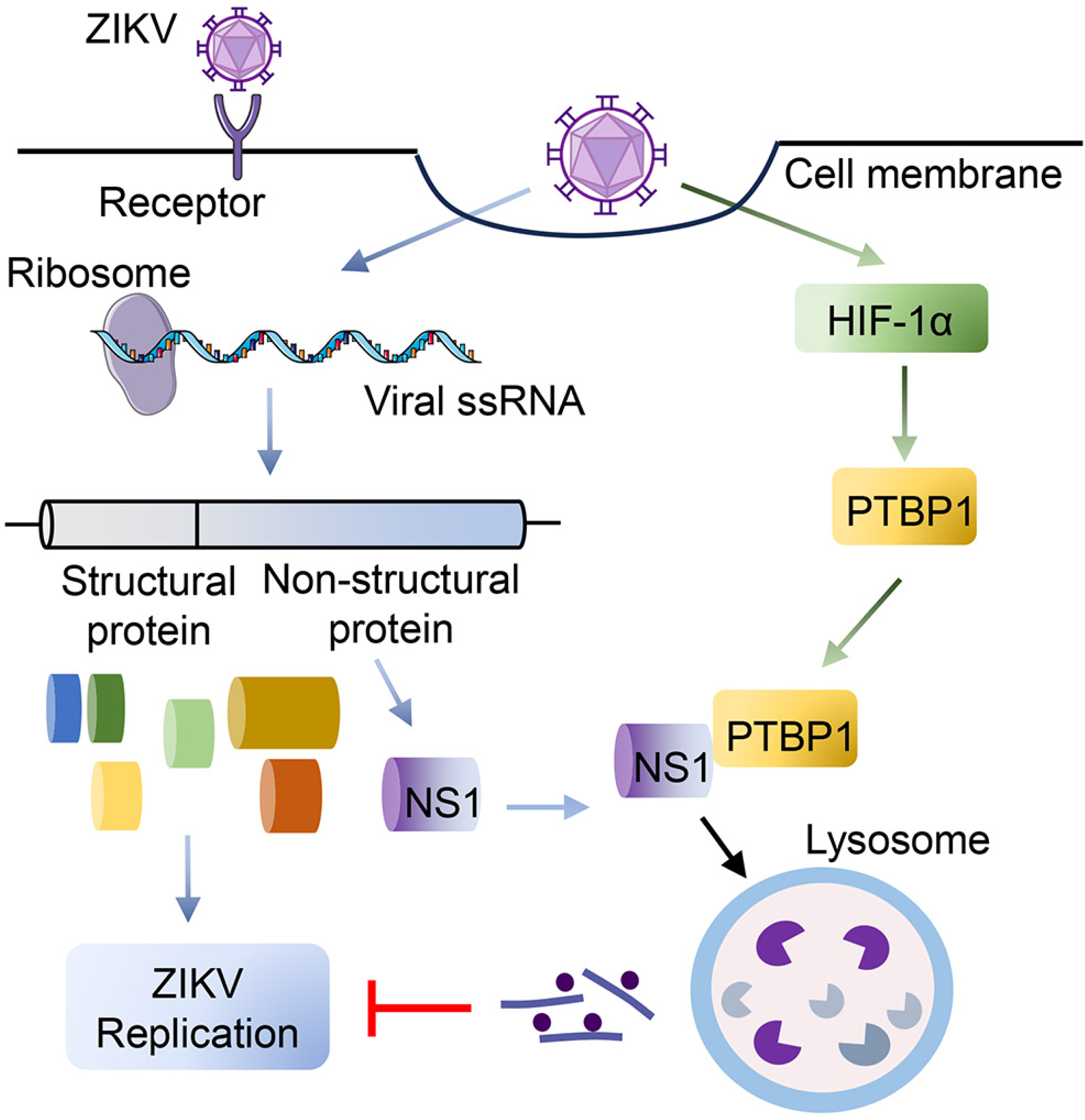
A proposal model underlying PTBP1 interacts with ZIKV NS1 to inhibit viral replication. Upon ZIKV infection, HIF-1α signal is activated to promote PTBP1 expression. Meanwhile, ZIKV viral RNA is translated into structural and nonstructural proteins, which participate in viral replication. The HIF-1α-induced PTBP1 interacts with ZIKV NS1 and then promotes the degradation of NS1 protein in a lysosomal pathway, and finally disrupts ZIKV replication.

## Discussion

ZIKV has previously triggered large-scale outbreaks globally, posing a severe threat to public health. However, to date, no effective vaccine or treatment is available. PTBP1 is an RNA-binding protein with a crucial role, and in recent years, its potential in inhibiting the replication of *Flaviviridae* viruses has gradually been unveiled. Specifically, Deepika et al. have shown that during Japanese Encephalitis Virus (JEV) infection, PTBP1 can translocate to the cytoplasm and effectively inhibit viral replication by competitively inhibiting the binding of JEV RNA to the NS5 protein (26). Jiang et al. have found that PTBP1 can bind to the NS4A protein and RNA of Dengue Virus (DENV), affecting virus production by regulating the synthesis of negative-strand RNA (27). The role of PTBP1 on ZIKV replication raises new insight into the interaction between PTBP1 and *Flaviviridae* viruses.

In our study, we observed that ZIKV infection significantly upregulates the expression level of PTBP1, a process that involves the activation of HIF-1α. In hypoxic environments, the expression of HIF-1α stabilizes and combines with HIF-1β to form HIFs, which regulate downstream signaling pathways and gene expression activity, thereby modulating various cellular adaptations and inflammatory responses (28, 29). Further studies indicate that modulating the expression level of PTBP1 directly restrains ZIKV replication. Specifically, HIF-1α induced-PTBP1 inhibits ZIKV replication, suggesting activation of HIF-1α inhibits viral replication. These findings further confirm the significant role of PTBP1 in the antiviral effect. In our previous finding, we reveal Severe Acute Respiratory Syndrome Coronavirus 2 (SARS-CoV-2) infection induces HIF-1α expression to aggravate inflammatory responses to Coronavirus Disease 2019 (COVID-19) (30). Notably, HIF-1α activation promotes SARS-CoV-2 replication, which differ this study that HIF-1α activation inhibits ZIKV replication by induction of PTBP1. As such, the control of HIF signaling is thought as a promising prospect for therapeutics in infectious diseases (31), including ZIKV infection and associated disorders.

Numerous studies have revealed that the interaction between host proteins and the non-structural proteins of ZIKV is one of the key mechanisms for inhibiting viral infection. For instance, the interferon-induced protein Viperin can specifically target the ZIKV NS3 and induce its degradation through the proteasome pathway, thereby effectively inhibiting viral replication (32). Similarly, the splicing factor Splicing factor 3b subunit 3 (SF3B3) protein can bind to the ZIKV NS5 and restrict viral replication by regulating GTP cyclohydroxylase1 (GCH1) (33). Here, we explore the specific mechanism by which PTBP1 exerts its antiviral effect. Furthermore, PTBP1 can induce the degradation of ZIKV NS1. We validated the specific binding of PTBP1 and ZIKV NS1 using Co-IP and immunofluorescence approaches, and determined their primary localization in the cytoplasm. PTBP1 interacts with the ZIKV NS1 to inhibit viral replication. The discovery of this new manner provides important theoretical support for the crucial role of PTBP1 in the antiviral process of ZIKV.

In eukaryotic cells, protein degradation is primarily accomplished through the ubiquitin-proteasome system and the autophagy-lysosome pathway, both of which play vital roles in maintaining intracellular homeostasis and protein quality control (34, 35). Previous studies have enlightened us with some clues. For example, ZIKV NS1 and ZIKV NS3 proteins have been found to bind to Poly (ADP-Ribose) Polymerase Family Member 12 (PARP12) and are degraded via the ubiquitin-proteasome pathway (36). In addition, C19orf66 has been shown to induce the degradation of ZIKV NS3 through a lysosome-dependent pathway (37). These findings suggest that PTBP1 may induce the degradation of ZIKV NS1 through a similar mechanism. To elucidate this issue, specific inhibitors are used to block the ubiquitin-proteasome system or the autophagy-lysosome pathway, respectively, and we observed that PTBP1-induced degradation of ZIKV NS1 is dependent on the lysosome pathway. This reveals a novel mechanism by which PTBP1 inhibits viral replication by specifically binding to and inducing the degradation of ZIKV NS1. These discoveries provide important insights into the understanding of the ZIKV replication process and the development of new therapeutic strategies.

In conclusion, this study reveals that ZIKV infection upregulates the expression level of PTBP1 by activating HIF-1α. PTBP1 exhibits a significant inhibitory effect on the replication process of ZIKV. In addition, PTBP1 can interact with ZIKV NS1 protein and induce its degradation through the lysosomal pathway, thereby inhibiting viral replication. This discovery provides a potential target for viral replication and pathogenesis in ZIKV-associated diseases.

## Acknowledgments

This work was supported by the National Natural Science Foundation of China [32400132].

## Author contributions

**Menglan Rao:** Conceptualization, Data Curation, Investigation, Methodology, Validation, Writing-original draft. **Zhiwei Lei:** Data Curation, Formal Analysis, Funding Acquisition, Investigation, Methodology, Validation. **Shuang Liu:** Data Curation, Formal Analysis, Investigation, Methodology. **Jiuxiu Lin:** Formal Analysis, Methodology, Resources. **Yue Kong:** Formal Analysis, Investigation, Methodology, Visualization. **Yicong Liang:** Formal Analysis, Methodology, Resources. **Zhen Luo:** Conceptualization, Project administration, Supervision, Validation, Writing-Review & Editing.

## Data Availability

All relevant data are available from the corresponding author upon request.

